# The ancestry of Antennapedia-like homeobox genes

**DOI:** 10.1101/2023.03.14.532566

**Authors:** Richard R. Copley

**Affiliations:** Laboratoire de Biologie du Développement de Villefranche-sur-mer (LBDV), Sorbonne Université, CNRS, 06230 Villefranche-sur-mer, France

## Abstract

I show that broad evolutionarily significant sub-groups of Antennapedia-like homeobox genes can be distinguished by consideration of 4 amino acids. The absence of proline at position 26 is a synapomorphy of the class; HOX-like homeodomains contain a 19,30 salt-bridge of inverted polarity with respect to the same residues in typical NK-like homeodomains, and residue 28 is highly conserved within but not necessarily between orthologous groups. None of these residues has an obvious role in sequence specific DNA recognition. The EH1 and hexapeptide sequence motifs outside the homeodomain are not well correlated with sub-type. From the discovery of a hexapeptide motif in a sponge NK-like gene, and identification of new instances of longer engrailed-like (EH2) variants of the hexapeptide, I infer that scattered motif distribution is unlikely to be due to convergent evolution, but rather multiple independent loss events. I reconcile these features and the species distribution of current genes to propose a scheme for the ordering of duplication events in the cnidarian-bilaterian stem group.

## Introduction

The homeobox was first identified as a part of genes in the Antennapedia and bithorax complexes, patterning the anterior-posterior axis in *Drosophila* [1–3], and homologous genes were found more or less straightaway in vertebrates and other animals [4,5]. The potential significance of this was apparent from the beginning [6,7]. The Hox genes, as they became known, belong to a much larger family of homeobox containing genes which, though not all homeotic, encode transcription factors with generally important roles in animal development. Various classes of homeobox containing genes have been proposed, based on phylogenetic analysis of the amino acid sequence of the homeodomain as well as the co-occurrence of the homeodomain with other protein domains. The HOX-like (including Hox, extended Hox and ParaHox genes) genes belong to what is now called the ANTP (Antennapedia) class^1^ [8]. In this scheme, non-HOX-like genes in the ANTP class are assigned to the NK-like subclass, named after the NK homeobox genes of Nirenberg and Kim [9]. Although genomes of single celled eukaryotes include homeobox encoding genes, the ANTP class appears to be specific to the Metazoa [10,11]. Of the two sub-classes (HOX and NK), there is a *prima facie* case that the NK-like genes are older than the HOX-like: HOX-like genes are absent from sponges, but a clear NK-like gene cluster has been discovered in *Amphimedon queenslandica* [12].

True orthologs of Hox genes are found in cnidarians, but not in sponges or ctenophores [13]. Based on synteny arguments and block duplication models, some authors have suggested there is indirect evidence for Hox loss in sponges, but this is contested [13,14]. A characteristic feature of the majority of Hox genes is the occurrence N-terminal to the homeobox of what is usually referred to as a ‘hexapeptide’ motif [15], although it generally only has 4 well conserved residues of consensus **YPWM** [16]. There are also a small number of NK-like ANTP class homeobox proteins with a hexapeptide-like sequence. This hexapeptide motif is the structural support for interaction with the TALE-class homeodomain proteins, enabling cooperative DNA binding [17]. The association of hexapeptide motifs with the homeobox has been shown to be functional in Cnidaria, but there is no evidence for its presence in sponges or ctenophores [18,19].

Another motif often found in homeobox containing proteins is the Engrailed Homology 1 (EH1) motif, of consensus FxSxxIL, which, in contrast to the hexapeptide, is most strongly associated with NK-like homeobox containing proteins, but also found encoded in several HOX-like genes, the parahox gene GSX and various Paired-class (PRD) homeobox genes [20]. This distribution led Bürglin and Affolter to suggest that the EH1 motif was present in the common ancestor of ANTP and PRD homeobox genes [21]. The EH1 motif mediates interactions with the transcriptional co-repressor Groucho/TLE. The engrailed genes, which give their name to the EH1 motif also encode an EH2 motif, described as “somewhat reminiscent of the hexapeptide” [15] or of “rudimentary similarity” [21], but equivalenced to the hexapeptide of Hox proteins by some authors. Sequence similarity between the two motifs is essentially limited to a single ‘conserved’ tryptophan: there is no YPWM core, even though the region of conservation in orthologs of engrailed is considerably longer than typical hexapeptides [22,23]. Like the hexapeptide, however, the region interacts with PBX1, a TALE-class homeodomain [22].

The evolution of ANTP homeoboxes has been extensively discussed within the context of three genomic clusters: the Hox cluster, which includes the Hox genes, several Hox-like genes (in human EVX, MNX, EN, GBX, MEOX) and DLX; the NK cluster and the Parahox cluster (see e.g. [24,25]. It has been hypothesised that the HOX and NK clusters were in turn linked into a single ‘mega-cluster’. Testing scenarios of evolution within or between these clusters has been limited as the short length of the homeodomain (60 amino acids) and generally high level of sequence conservation leading to a lack of phylogenetic resolution in the relative placement of orthologous families. To enquire further into the nature of the ancestral ANTP homeobox gene, I have examined protein sequence and structural motifs and gene order in a range of Metazoa, with special reference to non-bilaterians. I identify a clear, likely synapomorphic, structural delineation between a slightly redefined HOX-like class and other ANTP homeoboxes. I conjecture that the NK-class can be divided into ‘monophyletic’ NK genes (moNK) and a plesiomorphic subclass that likely gave rise to both the moNK genes and the HOX-like genes. I argue that the ancestral ANTP homeobox gene encoded both an EH1 motif and a hexapeptide-like sequence.

## Results and Discussion

### Comments on general scope

I collected alignments of orthologous sequences that were conserved in Bilateria (which I define as presence in both protostomes and deuterostomes - see methods). When cnidarian sequences were orthologous, I included these. All sequences and orthology group assignments are available at https://github.com/rcply/antp/. I have not included ‘orphan’ sequences, that is, isolated or likely misplaced long-branched sequences that did not show clear orthology with other bilaterians or cnidarians. Throughout I use a standard homeobox numbering of scheme of 60 amino acids where residue 50 is the Q of WFQN in the DNA recognition helix [4].

Most gene names are taken from their human orthologs (without the numeric distinction between paralogs e.g. EMX rather than EMX1). Several key orthology groups, however, are not present in human and in these cases I have used *Drosophila* or amphioxus names (Rough, Lms/Nedx, Nk7, Abox, Msxlx). When specific amino acid types are referred to at a particular site in a particular ortholog, they are generally obviously ancestral states for that ortholog because of conservation within both protostomes and deuterostomes. In some instances, individual species may be in conflict with this - for example echinoderm HOX11/13e of Szabó and Ferrier (2018) can encode an autapomorphic reversion to proline at residue 26 without affecting the main arguments presented below [26].

### Differential structural characteristics of and in the ANTP class

Some studies have attempted to define signature amino acids diagnostic of particular homeodomain families. Galliot and co-workers, for instance, identified PRD-class homeoboxes with 5 out of 6 diagnostic residues from P26 D27 E32 R44 Q46 [27] and Fonseca and co-workers ANTP [HQ][IV][AKLT] at positions 44-46 and [IKTV][ITV]W[FY]QN[HQR]R[AMNTVY]K at positions 46-55 as ANTP-like (i.e. present in HOXL and NKL subclasses, but not PRD homeodomains) [28].

Following the class assignments of human genes by Holland and co-workers, I constructed a multiple sequence alignment of metazoan PRD-class and ANTP-class orthologous sequences (also including some ANTP class genes lost from the human lineage - see methods) and calculated the cumulative relative entropy (CRE) of all positions, a method that identifies residues best able to discriminate between subgroups [29]. The highest scoring residue in the PRD vs. ANTP analysis, by far, was P26 (Z: 6.91 - **Figure 1**). Further inspection of aligned human homeodomains showed that in non-ANTP-class homeodomains residue 26 is always proline - in LIM, POU, HNF, SINE, TALE, CUT, PROS, CERS and ZF classes (with the exception of the 3rd homeodomain in ZHX1). In contrast, in the ANTP class it is generally leucine, although valine, isoleucine and occasionally methionine are also present.

**Figure 1:**
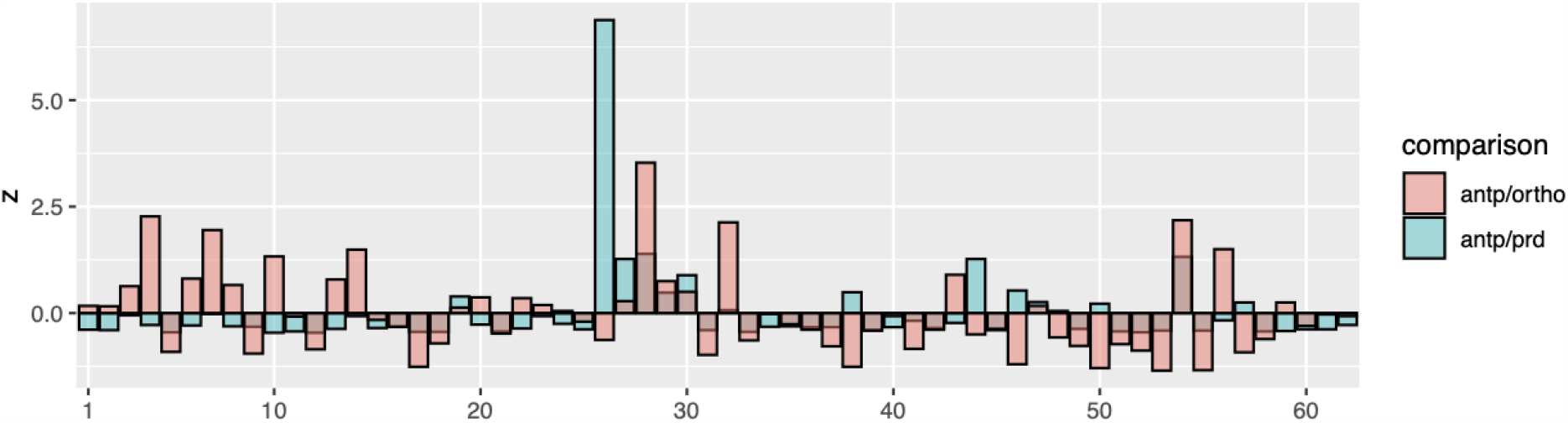
Cumulative relative entropy analysis. CRE Z-scores for PRD- vs. ANTP-class homeodomains (blue) and ANTP-class orthologous groups (pink). The x axis represents residue number in standard homeodomain numbering. For the PRD/ANTP comparison, residue 26 is the maximum, and for ANTP orthologs, residue 28.

Further analysis of CRE between orthologs of the ANTP class highlighted residue 28 (Z: 3.46) (**Figure 1**) which shows good conservation within orthologous groups, but often differs between them. Notably, in the HOX-like genes, that is, the Hox, Parahox, Evx, Mnx, Meox and Rough this residue is arginine. Exceptions to this include some posterior Hox genes (K28), Cdx (I28) and the Gbx orthology group (L28). No other ANTP orthology groups show arginine at this position. In contrast, for instance, Vax, Emx and Noto, together with Nk3 and Nk6 show a conserved glycine, and Engrailed a conserved glutamate (**see supplementary Table S1**).

In crystal structures, residue 28 sits above the phosphate backbone of the DNA molecule, but does not make base specific contacts [**Figure 2**]. R28 is required for full *ftz* activity in *Drosophila* [30,31], but the specificity of DNA binding can be accounted for without considering it [32,33]. Given the proximity to the phosphate DNA backbone, positively charged residues are readily explained, but outside the HOX-like genes, positively charged residues at this site are seldom observed. Alanine scanning experiments in Engrailed have shown a strong preference for alanine over the wild-type, negatively charged glutamate at position 28, and yet glutamate is conserved in all engrailed orthologs [34]. In the ANTP-class NANOG gene of vertebrates, L28A (L122A in NANOG numbering) increased protein stability and DNA binding strength [35]. The equivalent residue in the *Drosophila* paired gene (prd), isoleucine, is involved in dimer formation [36], but such a role is not readily apparent in any ANTP crystal structures.

**Figure 2:**
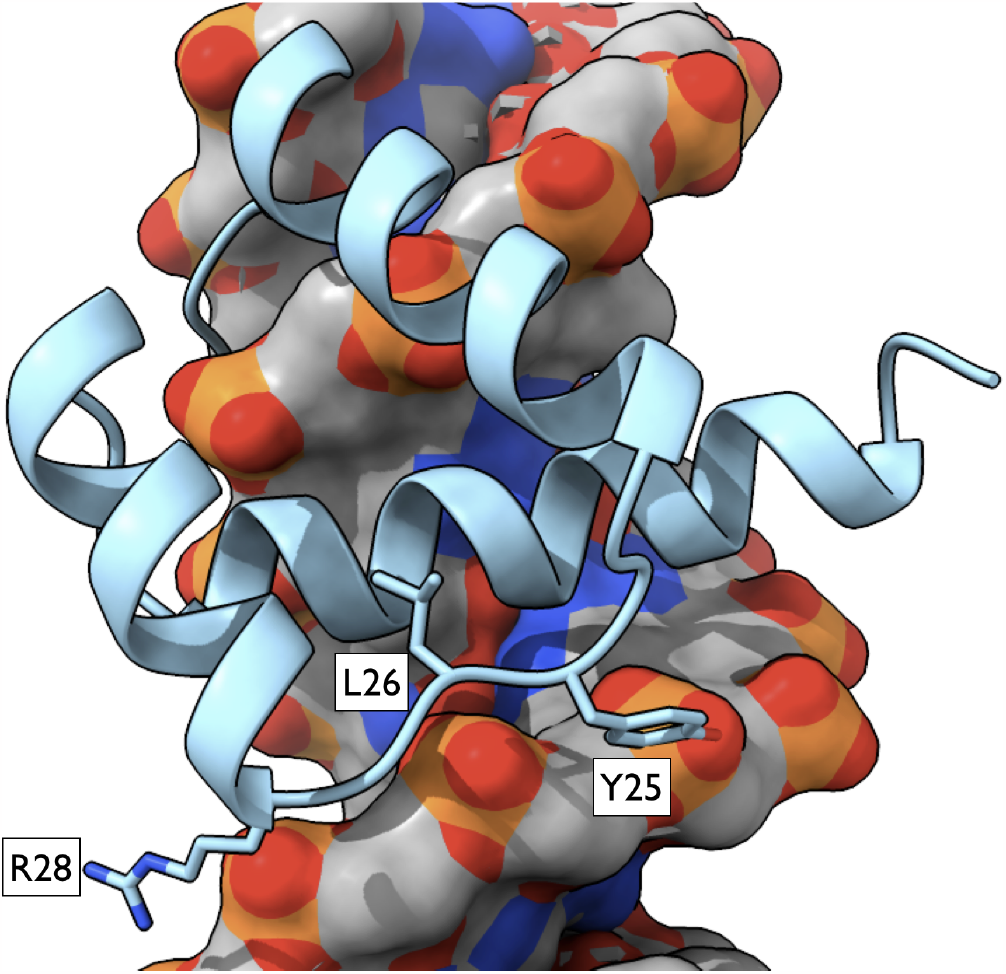
Orientation of the homeodomain bound to DNA. The C-terminal DNA recognition helix is towards the back of the protein sitting in the major groove of the DNA. The loop between helices 1 and 2, towards the bottom of the picture, tracks above the DNA sugar phosphate backbone. Y25, L26 and R28 are shown with sidechains (see discussion in text). Coordinates taken from the crystal structure of Antennapedia, RCSB PDB:9ANT.

In addition to the CRE analysis, I examined evolutionary couplings, that is, the correlations between residues in this alignment of ANTP-class sequences using the program ‘plmc’ [37]. The highest scoring pair of residues (19,30), involved in a salt bridge in some 3D structures, have previously been identified as highly correlated [38,39]. That the presence and polarity of these residue pairs has phylogenetic significance, however, has not been a focus of previous reports. In PRD-class genes, both residues are neutral. In HOX-like genes residue 19 is negative and residue 30 positive, whereas in NK-like genes residue 30 is always negative. In the majority of NK-genes residue 19 is positive, leading to the possibility of a salt bridge with inverted polarity relative to the HOX-like genes, although in several NK-like crystal structures residue 30 instead interacts with residue 23 (**Figure 3**). In other instances, (Barhl, Hmx, Dbx, Nk7, Nk6, Emx) residue 19 is uncharged, and in three cases (Gbx, Noto, Vax) both residues 19 and 30 are negatively charged (**Supplementary Table 1**). I conjecture that 19 -ve, 30 +ve is a synapomorphy of HOX-like genes (including all the true Hox and Parahox, Rough, Meox and Eve) and that this removes ambiguity in the classification of En (E19,R30) and Abox (E19,[RK]30), which become unequivocally HOX-like, as does Nedx/Lms. In contrast Dlx, another gene of disputed affinities [40] shows an (R19,E30) pair and so is NK-like. Two traditionally HOX-like genes are ambiguous in this scheme: Gbx (E19,E30) and Mnx (Q19,[KQ]30), although a potential *Trichoplax* ortholog of Mnx encodes the standard E19,R30 pair, suggesting a once HOX-like configuration.

**Figure 3:**
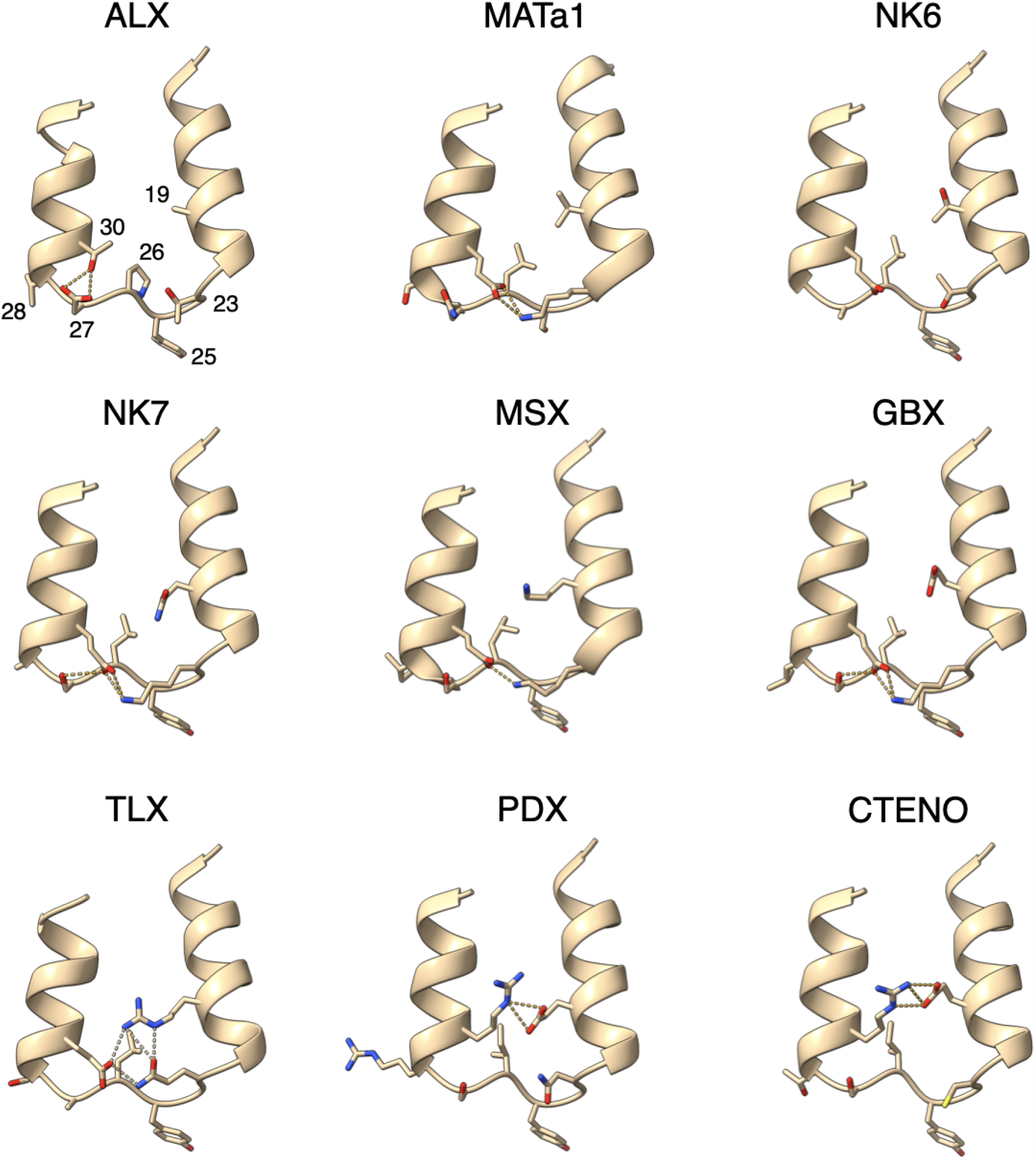
Varieties of hydrogen bond patterns around homeobox loop I-II. Hydrogen bonds are shown as dashed lines. Paired-like homeodomains (represented by ALX) show no salt bridge between residues 19 and 30 and a proline at residue 26. Using this same numbering scheme throughout, HOX-like ANTP (represented by PDX) domains have a negatively charged residue at position 19 and a positively charged residue at position 30. In NK-like homeoboxes, residue 30 is essentially always negatively charged. In typical NK-like domains (represented by TLX), residue 19 is positively charged, often forming a salt bridge with residue 30, of inverted polarity with respect to the HOX-like proteins, however, negatively charged residue 30 can also interact with residue 23 (MATa1, NK7, MSX, GBX). ANTP homeodomains have a non-proline aliphatic residue 26, usually leucine. Tyrosine 25 which hydrogen bonds to the DNA phosphate backbone is also shown (compare with **Figure 2** for orientation on DNA). N-terminal residues prior to helix I and post-helix II C-terminal residues including the DNA recognition helix have been removed for clarity. Red = oxygen, blue = nitrogen. MATa1 = a1 protein of *S. cerevisiae*. CTENO = *M. leidyi* ML102914a.

Aside from the 19,30 salt bridge, homeodomains often have two other salt bridges: between residues 17 and 52; and between residues 31 and 42 [39]. In neither case do they show an inverting polarity, although in Engrailed, residues 17 and 52 are both lysine.

### Sequence motifs outside the homeobox

As outlined in the introduction, ANTP class homeoboxes are often associated with either EH1 or hexapeptide motifs. I next searched for unreported instances of these, focussing especially on the more unusual associations of the hexapeptide in NK-like genes and the EH1 motif in HOX-like genes.

#### Hexapeptide motifs in putative TLX orthologs of homoscleromorph sponges

Relative to other sponges, Homoscleromorpha transcriptomes show good retention of genes shared with Bilateria [41]. I searched for homeobox homologs in the transcriptomes of the homoscleromorph sponges *Oscarella carmella, Plakina jani* and *Corticium candelabrum*. One homeobox containing gene, conserved in all three species, contained a hexapeptide motif (YPWM) perfectly matching the bilaterian consensus (**Figures 4 & S1**). The gene is a member of the NK-like subclass, with a +ve,-ve orientation of the 19,30 salt bridge, and also encodes an N-terminal EH1 motif. Its exact phylogenetic affinity is unstable in phylogenies, although it shows affinities with bilaterian TLX-like genes, which also encode an HX motif.

**Figure 4:**
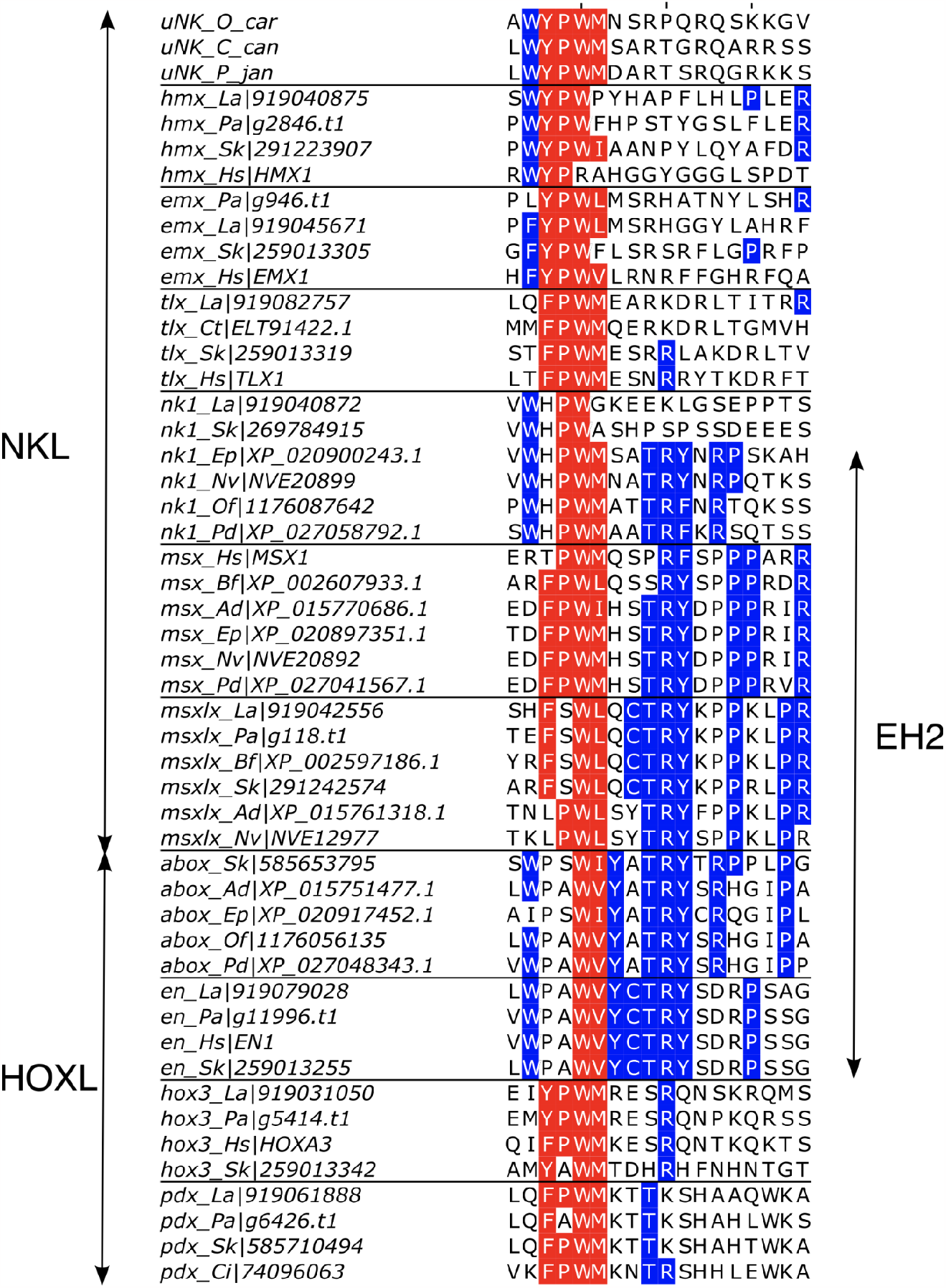
Alignment of selected hexapeptide and EH2-like motifs. The uNK sequences are from homoscleromorph sponges (see also **Figure S1**). Nk1, Msx, Msxlx, Abox and En share an extension of the hexapeptide. Classical hexapeptide residues are coloured red if they match the pattern W[YF]PW[MVIL]. Additional residues matching the EH2 consensus described here are coloured blue. Where possible, sequences are chosen to represent protostomes (La, Pa, Ct), deuterostomes (Hs, Sk, Ci) and cnidarians (Ad, Ep, Nv, Pd) - see methods for species abbreviations. Orthologous sequences showing no conservation of motifs have not been included.

Further transcriptome analyses revealed that all three sponges, but not non-homoscleromorph sponges, also encode orthologs of the PBX TALE class homeoboxes. PBX is the interaction partner of the HX motif in homeodomain proteins, and the sponge PBX sequences include the so-called PBC-A motif necessary for this interaction (**Figure S2**) [18].

#### Engrailed Homology 2 motifs in Abox, cnidarian MSX and NK1 class homeobox genes

Conservation between vertebrate and invertebrate engrailed genes led to the early definition of 5 ‘Engrailed Homology’ regions (EH1-5), with EH1 being the familiar Groucho interaction motif and EH4 being the homeodomain [20,22,42]. The EH2 motif (**Figure 4**) begins with a characteristic WPAW, the tryptophans of which were shown to be required for interaction with Exd/Pbx. The similar location (N-terminal to the homeodomain), conservation of the 2nd tryptophan and function (interaction with Exd/Pbx) led to the assumption of equivalence with the hox hexapeptide [22].

No convincing non-bilaterian orthologs of engrailed have been reported. While searching in cnidarians for genes containing EH2 motifs (as putative engrailed candidates), I identified genes with EH2 motifs in *Acropora digitifera* and other anthozoan cnidarians. Reciprocal searches and phylogenetic analysis suggested, however, that these sequences were orthologs of bilaterian Abox homeobox containing genes. The Abox gene (Absent in Olfactores) is not found in vertebrates or tunicates (i.e. Olfactores), but is present in other deuterostomes and protostomes [43]. Unlike bilaterian engrailed, orthologs conserve a salt bridge between residues E17 and R52 of the homeodomain. The Abox genes appear to have undergone further duplications in Anthozoa, and other cnidarian orthology groups do not encode an EH2-like motif. Intriguingly, however, these other putative Abox genes have a negatively charged residue, glutamate, at position 28, like bilaterian engrailed.

Further searching with an HMM constructed from representative sequences revealed clear examples of new EH2 motifs in cnidarian orthologs of Msx and Nk1 homeobox genes, as well as Msxlx genes. In general, these came from anthozoan cnidarians, which typically show slower rates of genomic evolution than medusozoans [44]. Like Abox, Msxlx is absent from vertebrates [45].

#### Continuity between EH2 motifs and hexapeptides, and potentially WRPW motifs

The EH2 motif as defined in Engrailed by Peltenburg and Murre extends over 18 amino acids. It can be seen (**Figure 4**), most clearly in cnidarian Msx sequences, that the motif includes an aligned hexapeptide-like motif anchored to this location by similarity in the more extended EH2 motif. In contrast, the EH2-defining Engrailed motifs show stronger conservation outside of the hexapeptide, in particular, in the ‘CTRY’ amino acid run, and a tryptophan immediately N-terminal to the hexapeptide itself. Taken together, these overlapping similarities provide direct alignment based evidence that hexapeptides and EH2 motifs are indeed homologous to each other, in agreement with others, although moving beyond a similar placement argument [23].

The homoscleromorph sponge hexapeptide motifs identified above are also immediately preceded by a tryptophan residue thus: **W**YPWM. Further, orthologs of the NK-like homeobox Hmx show a similar WYPWx motif N-terminal to their homeodomain, and various bilaterian orthologs of NK1 contain a conserved WxPW although they lack the remainder of the EH2 motif found in cnidarian NK1s.

NK like homeoboxes frequently contain an EH1 motif mediating their interaction with Groucho like co-repressors. A second motif known to interact with Groucho is found in hes-like bHLH transcription factors - its sequence, WRPW, is obviously similar to the WxPW EH2-submotif and hexapeptide motif I have highlighted here. Interestingly, WRPW motifs and YPWM motifs can exhibit similar backbone conformations, suggesting the possibility that they may be able to interact with common partners. Given the presence of a WYPWM motif in sponges, I speculate that the classical hexapeptide motif may have evolved from a motif that originally interacted with Groucho, and in proteins that still interact with Groucho, as revealed by the presence of additional EH1 motifs, the first tryptophan residue may be retained.

#### EH1 motifs in HOX-like genes

The EH1 motif is found in most NK-like homeodomain containing proteins and its presence has also been reported in HOX-like proteins including, for instance, Engrailed, Gbx and Mnx [20,46,47]. EH1 motifs are also found in orthologs of the HOX-like Nedx/Lms, Rough, Abox and Eve genes (**Figure 5**). In all cases the motif is towards the N-terminus of the protein. The Parahox gene Gsx also encodes a functional EH1 at its N-terminus [48]. This region is similar to the N-terminus of Hox9s and Hox10s (**Figure 5**). This latter similarity, but not the identification with the EH1 motif (or equivalent names), has been noted before [49]. These putative EH1 motifs overlap an ‘SSYF’ motif at the N-terminus of many Hox genes, shown to have a role in transcriptional activation [50], with the last two residues of the ‘SSYF’ being the first two of the EH1. Of note, these putative Hox and Parhox EH1s share a ‘DS’ residue pair with many EH1 representatives from NK-like proteins.

**Figure 5:**
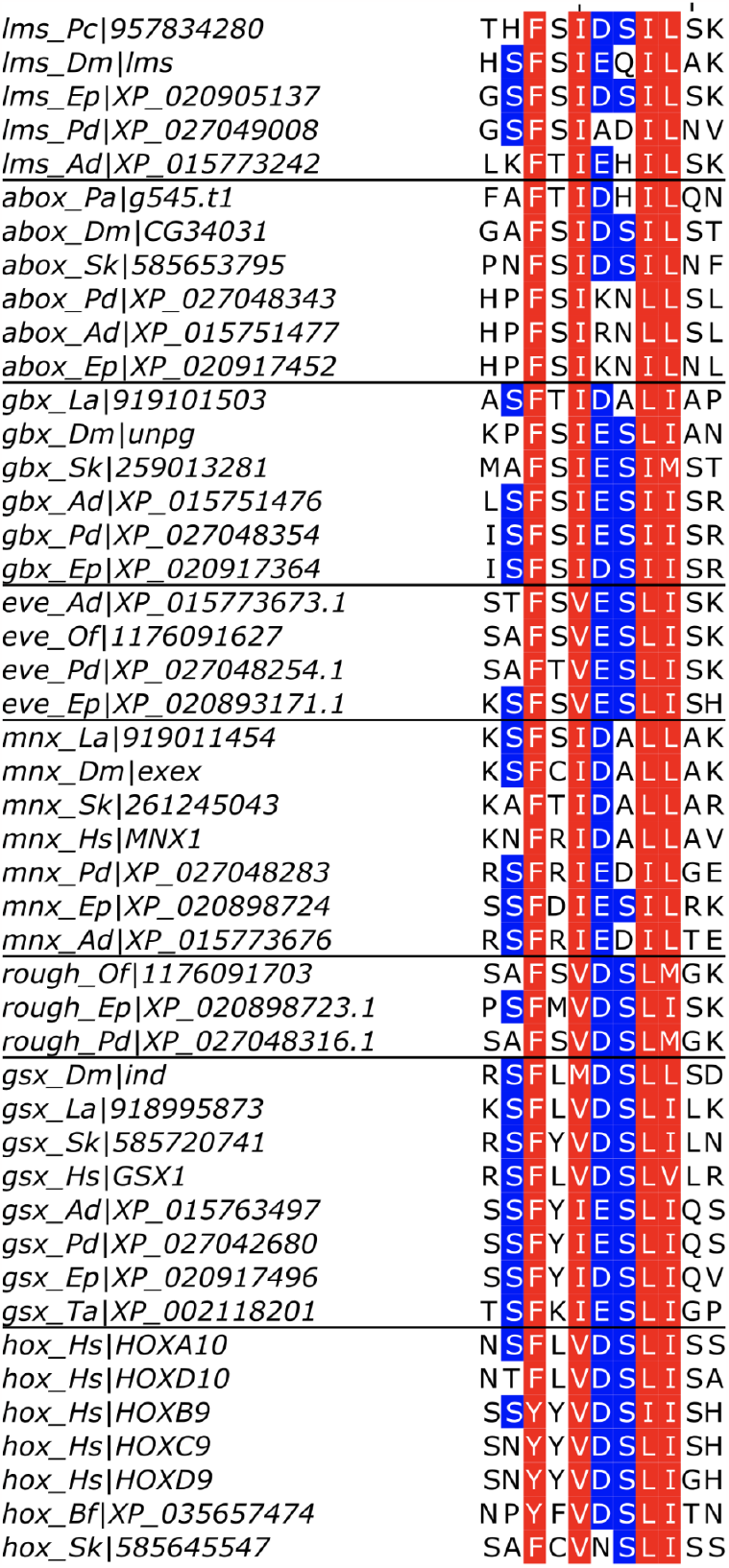
Alignment of selected EH1 motifs from HOX-like proteins. Residues matching the EH1 consensus [FY]x[IVM]xx[IL][IVLM] are coloured red. The central [DE]S pair found in Hox and the parahox GSX is coloured blue. See methods for species abbreviations.

### Genomic locations of EH2 containing homeobox genes

That the EH2 motif is found in a limited number of homeobox containing genes raises the possibility that the genes where it is found are more closely related to each other than to other homeobox containing genes. Relationships between different orthologous homeobox genes have been difficult to resolve, owing to their sequence conservation and short length, and studies have been informed by analysis of which genomic clusters genes belong to.

EN (engrailed) is a member of the so-called EHGbox cluster that, as originally defined in humans, also includes MNX (also known as HB9) and GBX [51]. ABOX (Absent in Olfactores) genes are not present in vertebrates, but are adjacent to EN in the hemichordate *Saccoglossus* and the sea urchin *Lytechinus*. In *Branchiostoma*, EN is adjacent to NEDX (Next to Distalless, known as LMS Lateral Muscles Scarcer in *Drosophila*, but again, missing in vertebrates), which is in turn adjacent to DLX **(Figure 6)**. In some cnidarians the ABOX gene is adjacent to GBX. In the cnidarian/bilaterian ancestor then, there is evidence of an EN, ABOX, GBX, MNX, NEDX, DLX cluster, although with EN/ABOX represented by a single ancestral gene. The ABOX/GBX pair are within 1.5Mb of a cluster of NK genes in *Acropora millepora* (NK1, HMX, HHEX and NK7) adding support for the idea of an ancestral ‘mega-homeobox’ cluster ancestral state of linked HOX-like and NK-like genes [52].

**Figure 6:**
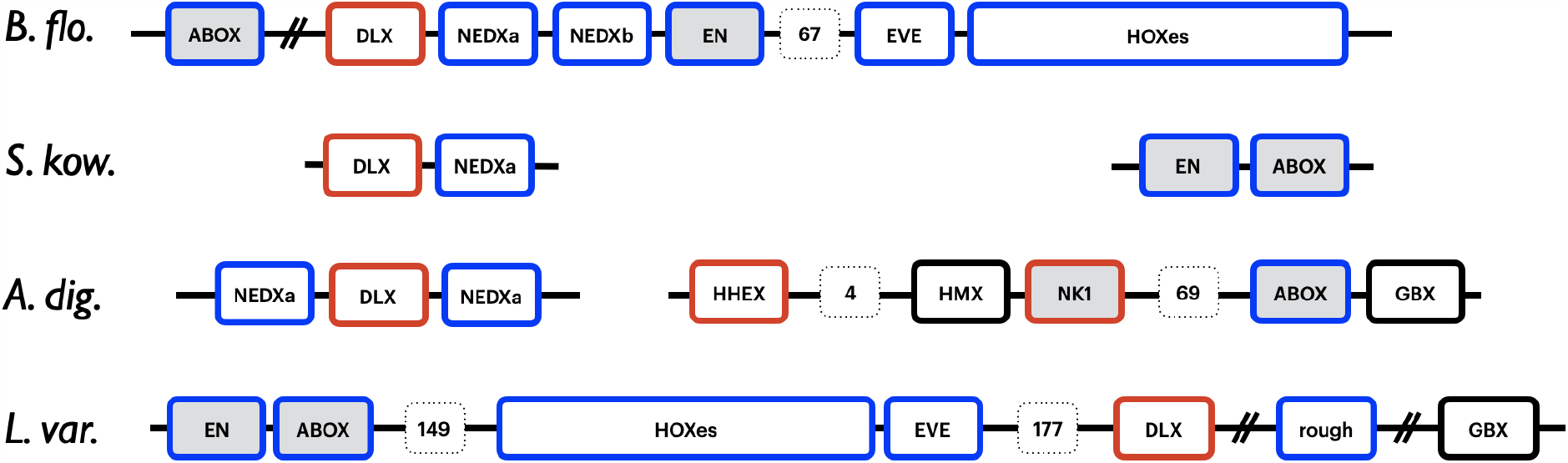
Genomic arrangement of key homeobox genes. 19-30 salt bridge polarity is indicated by the border colour: red border = NK-like; blue border = HOX-like. Boxes shaded grey indicate that the gene encodes an EH2 motif. Note that the ABOX and GBX genes are co-chromosomal with NK-like genes in Cnidaria and HOX-like genes in Bilateria. B. flo. = *Branchiostoma floridae*; S. kow = *Saccoglossus kowalevskii*; A. dig. = *Acropora digitifera*; L. var. = *Lytechinus variegatus*.

NK1 and MSX are canonical NK-class ANTP homeoboxes belonging to the NK cluster, (which also includes the hexapeptide containing TLX and HMX). The MSXLX gene is a member of the so-called ‘pharyngeal cluster’, adjacent to NK2-1 and NK2-2, that is conserved between *Trichoplax* and deuterostomes [41]. Given the similarity between MSX and MSXLX and that between NK2-1/2 and other NK genes, it is likely that this pharyngeal cluster is related to the NK cluster by a block duplication. A final non-hox hexapeptide encoding homeobox gene, EMX is clustered with VAX and NOTO, at least in humans - these may represent an extension of the NK-cluster.

In summary, the long variant of the hexapeptide, the EH2 motif, is present in the NK (MSX, NK1), the pharyngeal (MSXLX) and the EHGbox (EN, ABOX) homeobox clusters. The EHGbox cluster is linked to the ‘true’ hox genes; two of its constituents (MNX and GBX) are regarded as “HOX-like” on the basis of phylogenetic analysis, with EN close to the division between the NK-like and HOX-like subclasses, but classified as NK-like by Holland *et al*., although on the basis of linkage considered as an ‘extended-hox’ gene by others [18]. Taken together, these observations suggest that the EH2-like motif, which is longer than the hexapeptide and so unlikely to have arisen by convergence, was present in a common ancestral gene of all three clusters.

### Synapomorphies of HOX-like and NK-like homeoboxes

The NK-like homeoboxes that co-occur with an EH2 domain, NK1, MSX and MSXLX show the (R19,[ED]30) pairing. The HOX-like homeoboxes with an EH2 domain, EN and ABOX, show the (E19,[RK]30) pairing. The electrostatic nature of this residue pairing presumably limits the ease with which they can be interconverted, as to do so would require either an intermediate where both residues had the same charge, or via an additional step with a neutral residue (see later discussion). Aside from requiring multiple mutations, all extant sequences that could serve as potential intermediates between the EH2-containing NK-like genes and EN/ABOX (those with E19,E30, namely GBX, VAX and NOTO) lack EH2 motifs.

If this line of reasoning is correct, it suggests that all HOX-like and the (+ve19,-ve30) NK-like genes are mutually monophyletic with respect to each other. As such, a block duplication including a gene with an EH2 motif and a non-EH2 motif gene could not explain the relationship between a HOX-like and NK-like block, as it would, in addition, require multiple salt bridge inversions. This means instead that, barring convergent evolution, an EH2-like motif would have been present in the common ancestor of all HOX-like genes, and further, the common ancestor of (+ve19,-ve30) NK-like and HOX-like genes.

### On the possibility of sponge and ctenophore HOX-like sequences

The polarity of the 19,30 salt bridge (R19,[DE]30) implies that the hexapeptide containing homoscleromorph sponge sequences identified above are NK-class genes, in agreement with phylogenetic analysis. The sponge “CDX” gene identified by Fortunato *et al*., in contrast to bilaterian CDX does not contain a hexapeptide like motif [14]. The YIT motif presented by Fortunato *et al*. as another defining feature of CDX homeobox genes is also present in several other ANTP class homeoboxes (notably Rough, which is HOX-like) from various species. Further, the homeodomain itself lacks a small N-terminal extension diagnostic of bilaterian CDX homeodomains [53]. Although clearly of the ANTP class (with isoleucine, not proline at position 26), with E19,E30 and a lysine at residue 28, it is not CDX-like within the classification proposed here, but could represent a plesiomorphic HOX-like state similar to GBX (or NOTO/VAX).

Three closely related ANTP homeoboxes from the ctenophore *Mnemiopsis leidyi*, and an orthologous *Beroe forskalii* gene, show a HOX-like salt bridge polarity (**Figure 3 and S3**). These genes show no evidence of encoding a hexapeptide and their relationship to specific orthologous groups within the ANTP class cannot be ascertained, but it may prove relevant that some members also include a YIT motif as in the sponge “CDX” above (**Figure S3**). No ctenophore homeodomain sequences contain an obvious hexapeptide sequence motif. One *Mnemiopsis* sequence (ML02215a) does contain a pair of tryptophans two residues apart, N-terminal to the homeodomain (**WY**S**W**V), in a manner reminiscent of the N-terminal half of the EH2 motif. This motif is also present in the very similar *Beroe forskalii* ortholog (**Figure S4**). These homeodomains do not show obvious affinities to any specific bilaterian ANTP orthology group, but show an NK-like 19,30 salt-bridge (R19,E30).

### A scenario

Although homeodomain phylogenies are typically poorly resolved, studies have consistently suggested that the PRD class is sister to the ANTP class e.g. [27,54,55], and notably, both classes share proteins with an EH1 motif N-terminal to a homeodomain. EH1 motifs in PRD class homeodomain proteins often start with histidine instead of the more common phenylalanine [20]. The only ANTP-class genes where this is the case are the NK6-like. Of the features identified in this study, no proteins include both a PRD-like P26 residue and an ANTP-like salt bridge between residues 19 and 30. A few ANTP-like proteins, however, lack any sort of hydrogen bonding possibility between helices I and II, and this is close to a PRD-like state. NK6 is one of these proteins. I propose that the plesiomorphic condition of the ANTP class was similar to NK6, and that the salt-bridge containing ANTP proteins (that is, the HOX-like and monophyletic NK-like) are derived states. Examination of extant ANTP sequences and structures suggests a mechanism for apparent inversion of the polarity of the 19,30 salt bridge, while minimizing potentially deleterious electrostatic interactions via stabilizing charges on residue 23 (**Figure 7**). A negatively charged residue at position 30 acts as an acceptor for hydrogen donors in the I-II loop (this is also seen in some TALE class homeodomains). This residue can form salt bridges with charged residues at positions 19 or 23 in different NK-like homeodomains (**Figure 3**). In the subclass of homeodomains that would become HOX-like, residue 19 became negatively charged (as in Gbx: -ve,-ve), and a subsequent mutation led to residue 30 becoming positively charged, inverting the polarity of the salt bridge (-ve,+ve) relative to the NK-like genes.

**Figure 7:**
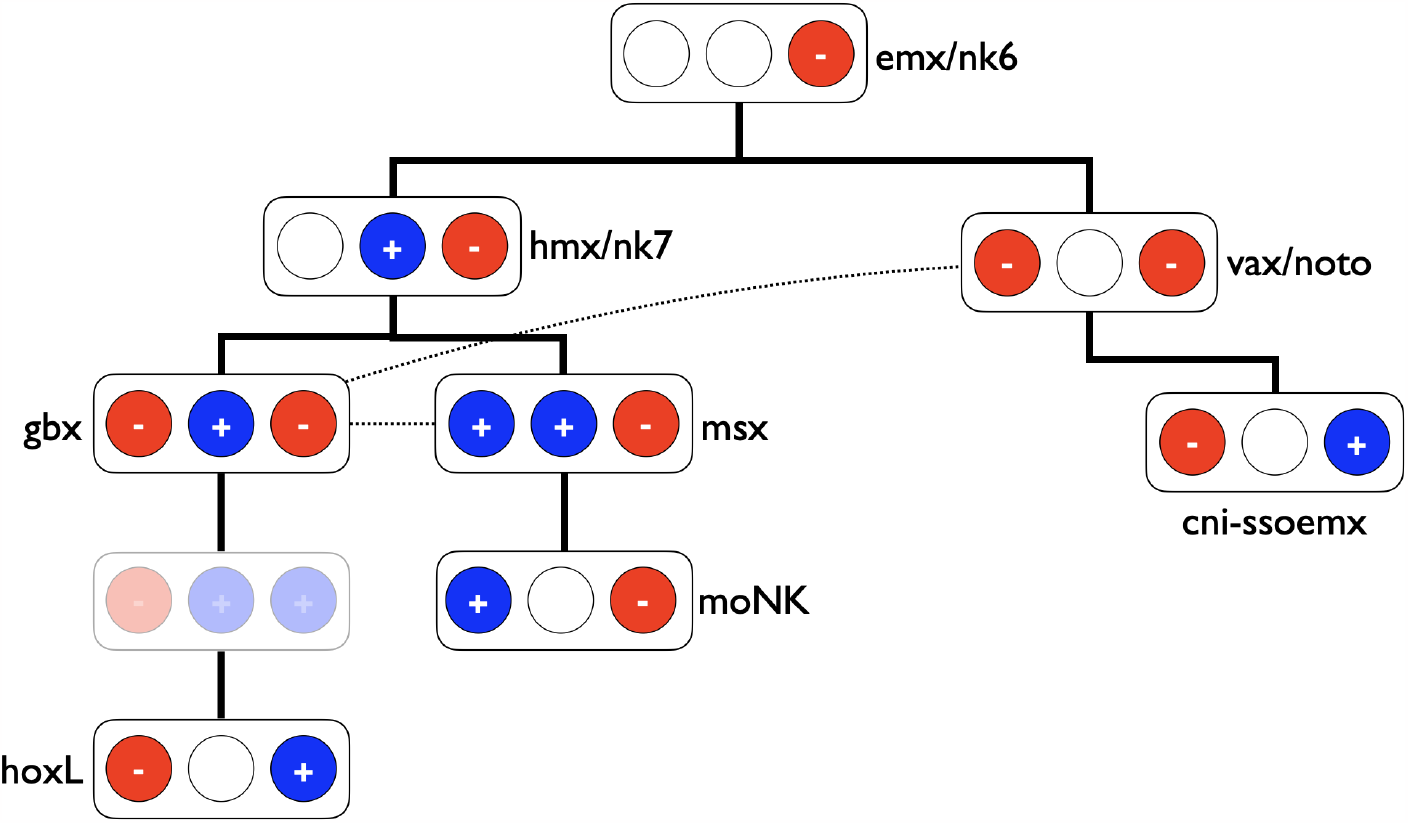
Charges of residues 19,23 & 30 in ANTP-like homeoboxes. Circles represent residues 19,23 & 30 from left to right (standard homeodomain numbering from 1-60; see text for details), with red negative, blue positive and white uncharged. States are arranged in a tree-like manner with one state change between ‘generations’, providing one plausible pathway for the salt-bridge polarity inversion between HOX-like and NK-like genes. Dotted lines show alternative routes. Representative gene names are given but should not be conflated with ancestral genes. hoxL = HOX-like; moNK = monophyletic-NK

Although these salt bridge configurations are compatible with gene clusters serving as phylogenetic proxies (i.e. NK, Hox and Parahox and pharyngeal clusters), it is apparent that the distribution of sequence motifs is not. EH1 and Hexapeptide motifs (or the variant EH2) are found in both NK- and HOX-like genes, although they are also absent in numerous representatives from the same likely clades. Taken together with the fact that I have identified a sponge gene that encodes EH1, Hexapeptide and Homeodomain, this suggests that the likely ancestral ANTP-like gene also encoded all three features. This is consistent with the placement of EMX and HMX as being similar to the ancestral ANTP-like state.

NK6 forms a clade with NK7 and HMX (aka NK5) genes, and representatives of this clade are found in sponges [12]. Similarly, EMX, VAX and NOTO consistently form clade under a variety of models, along with sponge representatives. Both EMX and HMX like genes encode EH1 and hexapeptide-like motifs in bilaterians (**Figure 4**). In the scenario outlined in **Figure 7** the GBX-lke state is a crucial intermediate in the evolution of HOX-like genes. GBX itself lacks an EH2 or hexapeptide motif, so taken at face value, this implies that the GBX/HOX lineage must have diversified rapidly, before the ur-GBX had an opportunity to lose its hexapeptide motif. Similar arguments may account for the loss of EH1 motifs at the N-terminal end of anterior HOX genes, but not, for instance, the parahox gene GSX.

The questions arise as to whether there is any biological significance to the loss of proline as a defining feature of ANTP-class homeodomains, and the significance of the salt bridge inversion between HOX and NK-like homeodomains - that is, do Hox genes only work as Hox genes at the organismal level because their salt bridge has this polarity? In the case of loss of proline, Fersht and co-workers have argued that the conformational flexibility of the network of residue interactions centred around L26 is important for induced fit interactions with DNA [56]. Salt bridge polarity can have subtle but unpredictable effects on function - in GBX, inverting the polarity of the 17,52 salt bridge causes a loss of DNA binding affinity through transient molecular interactions [57]. At a similarly subtle level, it is not clear how the precise nature of homeodomain residue 28 affects the biological roles of different ANTP-like homeodomains - and yet the evidence discussed above suggests that it does.

## Methods

### Identification of homeobox sequences, master alignment and phylogeny

I used hmmer to search a representative set of metazoan proteins derived from complete genome sequences with the Pfam Homeobox (PF00046.28) Hidden Markov Model, selecting hits based on the ‘gathering’ threshold (‘--cut_ga’) [58]. Sequences were filtered and aligned as described in [41]. I inferred a phylogenetic tree using iqtree with the LG+C20 model.

#### Core species searched

Cnidarians: Ad, *Acropora digitifera*; A_aur, *Aurelia aurita*; C_hem, *Clytia hemisphaerica*; D_gig, *Dendronephthya gigante*; Ep, *Exaiptasia pallid*a; Hv, *Hydra vulgaris*; M_vir, *Morbakka virulenta*;Nv, *Nematostella vectensis*; Of, *Orbicella faveolata*; Pd, *Pocillopora damicornis*;

#### Deuterostomes

Hs, *Homo sapiens*; Ci, *Ciona intestinalis*; Bf, *Branchiostoma floridae*; Sk *Saccoglossus kowalevskii*; Sp *Strongylocentrotus purpuratus*; Xb, *Xenoturbella bocki*; P_nak, *Praesagittifera naikaiensis;*

#### Protostomes

Ac, *Aplysia californica*; Ce, *Caenorhabditis elegan*s; Cg, *Crassostrea gigas*; Ct, *Capitella telata*; Dm, *Drosophila melanogaster*; La, *Lingula anatina*; Pa, *Phoronis australensis*; Pc, *Priapulus caudatus*; Homoscleromorph sponge transcriptome assembly was described previously [41].

### Construction of ANTP, PRD and orthology group subalignments

ANTP and PRD regions of the complete homeobox phylogenetic tree were inspected for orthologous groups, including protostome and deuterostome (vertebrate + ambulacrarian) sequences, corresponding to accepted ANTP and PRD human homeobox families [8], adding in conserved families that have been lost in humans (Antp class: Abox, Bari, Msxlx, Nedx/lms, Nk7, Rough).

Where cnidarian sequences were available, these were included. For each family, I extracted a representative 62 amino acid sequence, such that residue 50 was the equivalent of the typical Q50 in most ANTP homeoboxes. These were then searched against the full length sequences of the ortholog group using glsearch3 from the FASTA package and ungapped 62 amino acid fragments (i.e. those reported with cigar string ‘62M’) extracted to make an ungapped multiple sequence alignment. The alignments were merged to form a single alignment with each sequence coming from a known orthology group. Protostome/Deuterostome orthology of central and posterior Hox genes is hard to establish, so I treated orthology groups 5-8 as a single central group (HOX-c), and 9-13 as a single posterior group (HOX-p).

Alignments are available at: https://github.com/rcply/antp

### Sub-type specific amino acids

Aspects of the methodology of PROUST (Prediction of Unknown Subtypes) to identify specificity determining residues [29] were reimplemented in python, and using Hidden Markov Models of the HMMER3 package [58]. Orthology or higher level evolutionary groupings (e.g. ANTP vs PRD) were used to define the groups for which the most likely specificity determining residues were to be found, and over which cumulative relative entropy was calculated.

Code is available at: https://github.com/rcply/antp

### Gene order

Gene locations were taken from GFF annotation files associated with genome builds. Gene identifiers were cross referenced with the homeobox Pfam Hidden Markov Model searches and orthology identification from the phylogenetic analysis.

### Protein structure

Where crystal structures were available, these were used as data in Figure 3. In the case of NK6, NK7 and the ctenophore sequence, locally run Alphafold2 predictions were used, with relaxation [59,60]. I also used an Alphafold2 prediction in preference to the available NMR structures of GBX. There can be little doubt that the global folds of these predictions are correct, and side chain locations with hydrogen bonding patterns appear highly plausible. Further, I have not drawn conclusions on absence of hydrogen bonding potential unless the sidechains of involved residues were incapable of this. RCSB accessions for crystal structures: ALX = 3a01_B; MATa1 = 1AKH_B; MSX = 1ig7_A; TLX = 3a01_A; PDX = 2h1k_A. Hydrogen bonds were calculated between all illustrated side chains using Chimerax (i.e. ‘hbonds (sel & sidechain) restrict both’).

## Acknowledgments

Thanks to Professors Graham Budd (Uppsala) and Max Telford (UCL).

## Supplementary Material

**Table S1:** Key homeodomain residue types. The most frequent residue at each position is shown (see supplementary data). Residues 19 & 30 have potential to form a salt bridge, and this is observed in many structures, with NK-like genes having a different polarity to HOX-like genes. Residue 23 can also form salt bridges with residue 30. Gene names prefixed with ‘cni-’ are cnidarian specific (see supplementary data). HOX-c and HOX-p are central and posterior type Hox genes.

|  | 19 | 23 | 28 | 30 |
| --- | --- | --- | --- | --- |
| GSX | E | N | R | R |
| PDX | E | N | R | R |
| CDX | E | S | I | R |
| cni-CDX | E | C | R | R |
| HOX1 | E | N | R | R |
| HOX2 | E | N | R | R |
| HOX3 | E | N | R | R |
| HOX4 | E | N | R | R |
| HOX-c | E | N | R | R |
| HOX-p | E | N | R | R |
| MEOX | E | N | R | R |
| MNX | Q | N | R | K |
| EVE | E | E | R | R |
| ROUGH | E | N | R | R |
| ABOX | E | D | V | K |
| LMS | E | N | V | R |
| EN | E | N | E | R |
| GBX | E | K | L | E |
| NK7 | Q | K | A | E |
| HMX | T | K | S | E |
| MSX | K | K | I | E |
| BARI | K | K | I | E |
| BARX | R | Q | T | D |
| BSX | R | Q | T | E |
| DBX | M | Q | K | D |
| HLX | R | Q | K | D |
| BARHL | S | Q | V | D |
| DLX | R | T | L | E |
| HEX | K | Q | P | E |
| NK1 | K | T | V | E |
| TLX | R | Q | S | E |
| LBX | R | Q | P | D |
| NK3 | R | Q | G | E |
| MSXLX | K | T | S | D |
| NK2-3/5/6 | R | Q | A | E |
| NK2-1/4 | R | Q | A | E |
| NK2-2/8 | R | Q | A | E |
| NK6 | T | T | G | E |
| EMX | A | N | G | E |
| NOTO | E | Q | G | E |
| VAX | E | N | G | D |
| cni-SOEMX | R | N | G | E |
| cni-SSOEMX | E | D | G | K |
| cni-SOVENTX | I | Q | R | E |

**Figure S1.**
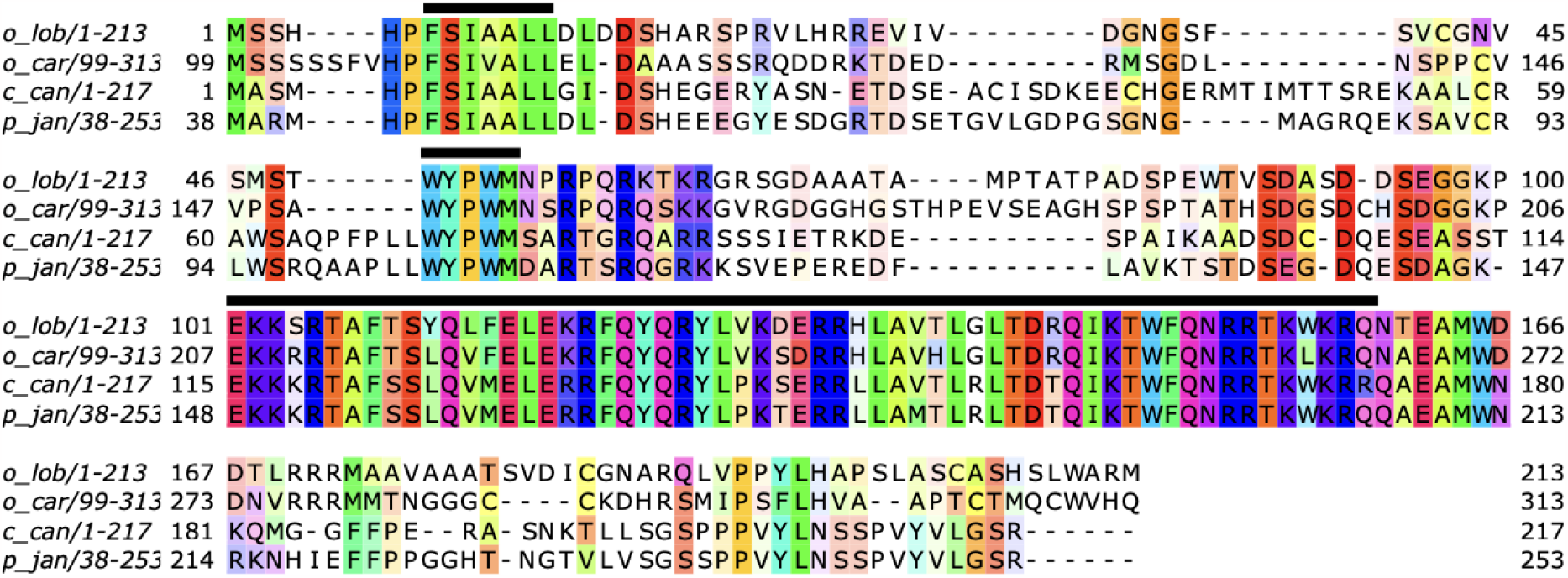
The homoscleromorph sponge genes with hexapeptides described in the text. From N- to C-EH1, hexapeptide and homeobox sequences are marked with black bars. o_lob = *Oscarella lobularis*; o_car = *Oscarella carmela*; c_can = *Corticium candelabrum*; p_jan = *Plakina jani*

**Figure S2.**
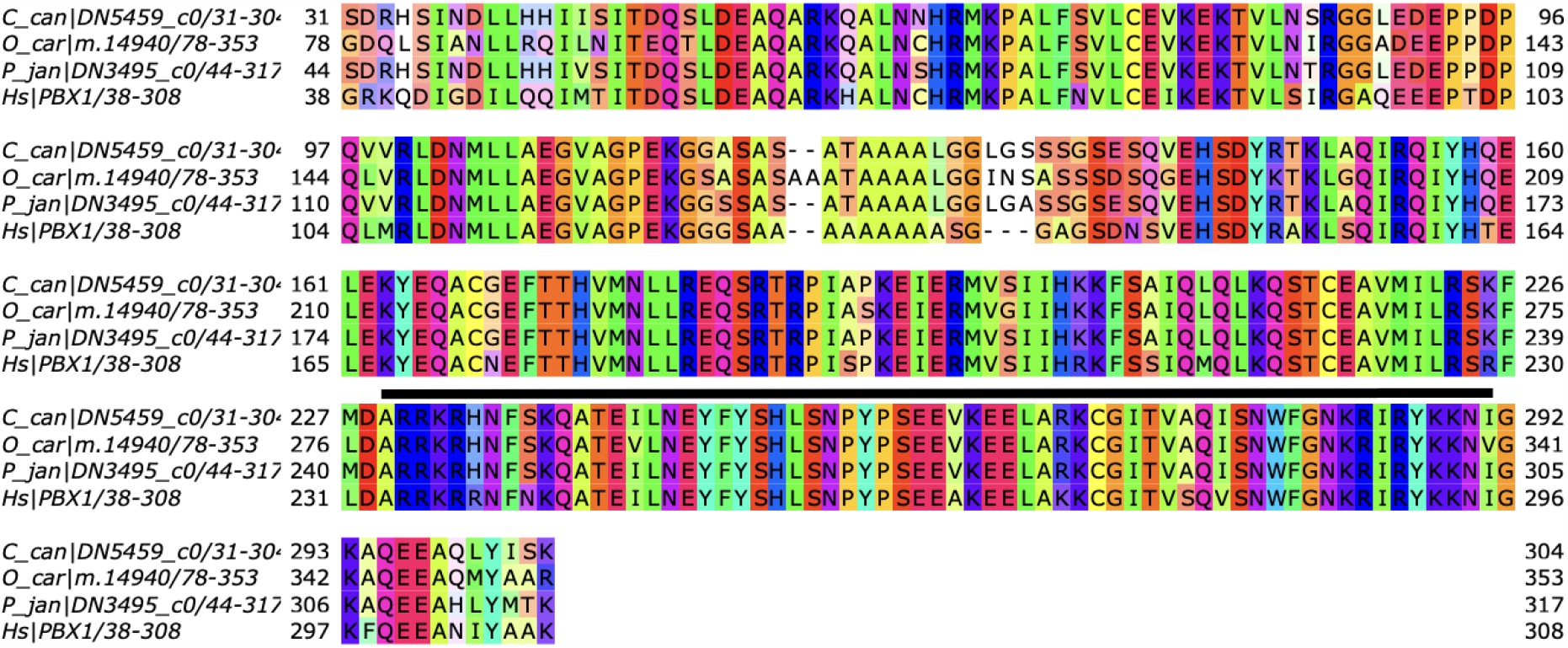
Homoscleromorph sponge orthologs of the TALE-class PBX. Alignment trimmed to show regions of maximum conservation with human PBX1. The black bar indicates the homeodomain. Sponge species as figure S1.

**Figure S3.**
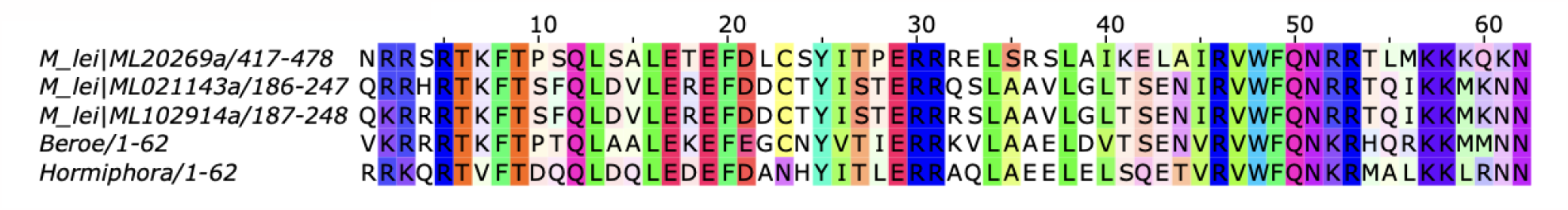
Ctenophore homeodomain sequences with potential for a HOX-like salt bridge polarity (-ve,+ve) between residues 19 & 30

**Figure S4.**
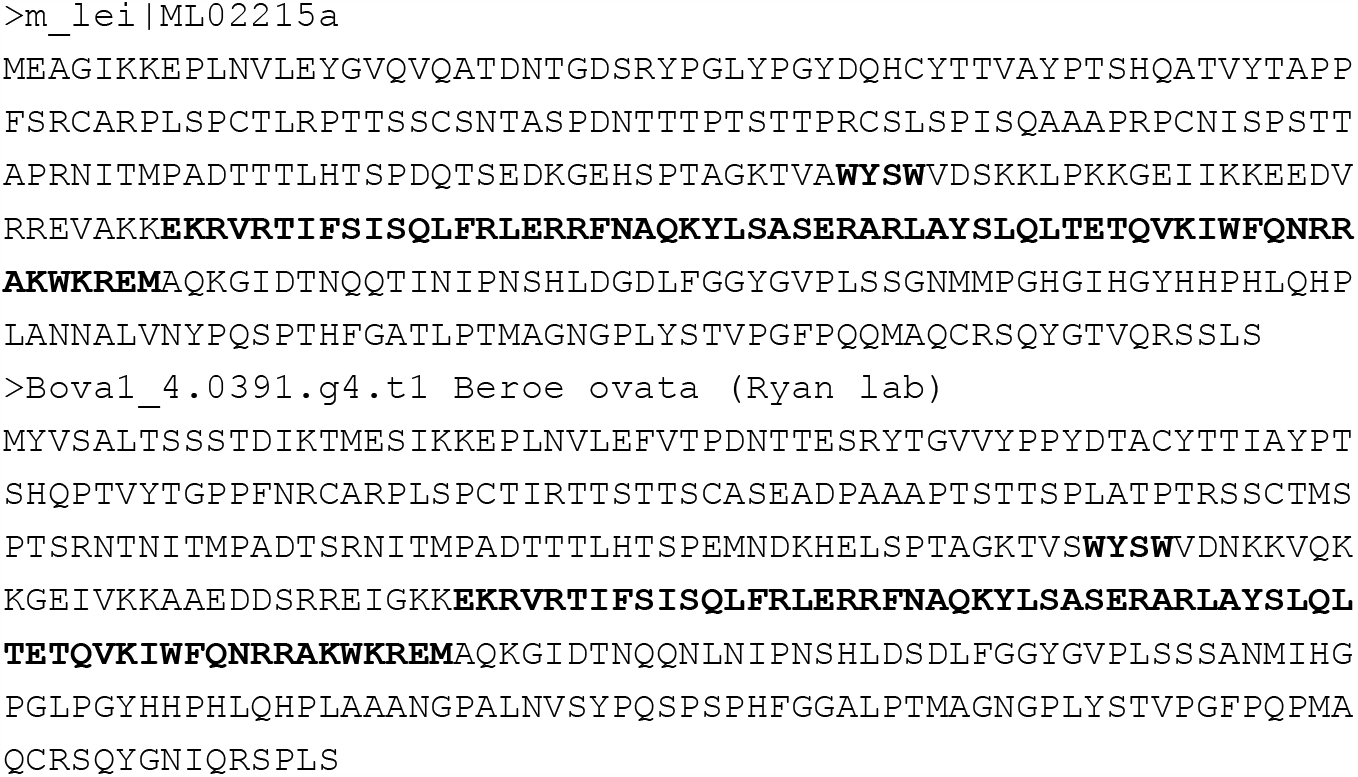
Ctenophore sequences with WYSW motif. Motif and homeobox in bold.

## Footnotes

1 Early use of ‘ANTP class’ is less well defined, but generally narrower and more loosely interpretable as sequence similarity to Antennapedia or ability to hybridise to particular probes.

